# Epithelioids: Self-sustaining 3D epithelial cultures to study long-term processes

**DOI:** 10.1101/2023.01.03.522589

**Authors:** Albert Herms, David Fernandez-Antoran, Maria P. Alcolea, Argyro Kalogeropoulou, Ujjwal Banerjee, Gabriel Piedrafita, Emilie Abby, Jose Antonio Valverde-Lopez, Inês S. Ferreira, Stefan C. Dentro, Swee Hoe Ong, Bartomeu Colom, Kasumi Murai, Charlotte King, Krishnaa Mahbubani, Kourosh Saeb-Parsy, Alan R Lowe, Moritz Gerstung, Philip H Jones

**Affiliations:** Wellcome Sanger Institute, Hinxton CB10 1SA, UK; Department of Biomedical Sciences, Faculty of Medicine, Universitat de Barcelona, Barcelona 08036, Spain; Cell Compartments and Signalling Group, Institut d’Investigacions Biomèdiques August Pi i Sunyer (IDIBAPS), Barcelona 08036, Spain; Wellcome/Cancer Research UK Gurdon Institute, University of Cambridge, Tennis Court Road, Cambridge CB2 1QN, United Kingdom; Wellcome-MRC Cambridge Stem Cell Institute, Jeffrey Cheah Biomedical Centre, Cambridge Biomedical Campus, University of Cambridge, Puddicombe Way, Cambridge, CB2 0AW, UK; Department of Oncology, University of Cambridge, Hutchison Research Centre, Cambridge Biomedical Campus, Cambridge CB2 0XZ, UK; Spanish National Cancer Research Centre (CNIO), Madrid 28029, Spain; Department of Biochemistry and Molecular Biology, Complutense University of Madrid, Madrid, Spain; European Molecular Biology Laboratory, European Bioinformatics Institute, Cambridge, UK. Present Address: Artificial Intelligence in Oncology (B450), Deutsches Krebsforschungszentrum Im Neuenheimer Feld 280, 69120 Heidelberg, Germany; Department of Surgery and Cambridge NIHR Biomedical Research Centre, Biomedical Campus, University of Cambridge, Cambridge CB2 2QQ, UK; Institute for Structural and Molecular Biology, University College London, London, United Kingdom; Institute for the Physics of Living Systems, University College London, London, United Kingdom; Department of Physics and Astronomy, University College London, London, United Kingdom

## Abstract

Studying long-term biological processes such as the colonization of aging epithelia by somatic mutant clones has been slowed by the lack of suitable culture systems. Here we describe epithelioids, a facile, cost-effective method of culturing multiple mouse and human epithelia. Esophageal epithelioids self-maintain without passaging for at least a year, recapitulating the 3D structure, cell dynamics, transcriptome, and genomic stability of the esophagus. Live imaging over 5 months showed epithelioids replicate *in vivo* cell dynamics. Epithelioids enable the study of cell competition and mutant selection in 3D epithelia, and how anti-cancer treatments modulate the competition between transformed and wild type cells. Epithelioids are a novel method with a wide range of applications in epithelial tissues, particularly the study of long term processes, that cannot be accessed using other culture models.

## Introduction

The long-term culture of primary epithelial cells has been challenging. A wide range of tissues may be expanded using organoid cultures^1-4^. These have been used to model epithelial carcinogenesis and genetic disease to guide therapy ^5-9^. However, organoids and similar models have a critical limitation. They continuously expand, and therefore do not attain the balance between proliferation and differentiation that maintains adult tissues in a steady state, and so require frequent passaging^10-15^. Classical organotypic cultures where epithelial cells are grown on permeable membrane inserts give excellent 3D epithelial histology but only last for a few weeks^16,17^. Older methods such as explant culture, where primary cells migrate out of tissue sample onto a plastic surface fail to replicate 3D tissue structure ^18^. None of these systems retains both the architecture and the long-term self-maintaining capacity of adult epithelia.

These limitations have restricted the study of processes that require long-term, self-maintaining, epithelial cultures. *In vivo*, mutant clones expand over months to years, to multi millimeter sizes competing for the limited space available in the proliferative compartment of homeostatic epithelia ^19-24^. Lineage tracing using transgenic mouse models can capture many of the features of mutant clonal dynamics in human epithelia but are slow and not suitable for screens to uncover the genes that regulate competitive fitness ^19,25-30^.

To overcome these limitations, we have developed ‘epithelioids’, suitable for studying processes that require long-term, centimeter scale epithelial cultures, by combining explant culture with organotypic protocols to generate 3D *ex vivo* epithelial reconstructions. This system avoids the stress of disaggregating tissue into a single cell suspension, allowing substantial *in vitro* tissue amplification, and uses a medium extensively tested for skin grafting in humans, at a lower cost than standard organoid protocols^3,15,31^. Moreover, we demonstrate that epithelioid cultures replicate tissue organization and gene expression and self-maintain for at least one year. This allows the study of long-term tissue and cellular processes in a 3D epithelial context.

## RESULTS

### Generation of epithelioid cultures from mouse and human epithelia

We began by culturing mouse esophageal epithelial explants on permeable membrane inserts in a standard keratinocyte medium supplemented with growth factors (cFAD, methods) (**Fig.1a**)^15^. Epithelial cells migrated out from the explant forming an expanding cellular ‘halo’ of keratinocytes within 48h, which grew at a constant velocity (0.52 mm^2^/h) until the cultures reached confluence (**Extended Fig.1a-c and Supplementary Video 1**). One week after plating, tissue explants can be removed. Cells from a single explant (of up to 2.5 mm^2^ or 1/32th of the esophagus) covered a 4.5 cm^2^ membrane in 15-18 days, a 182 fold increase in area. Establishing cultures was highly efficient, with 95% of explants generating keratinocyte outgrowth (from multiple mice and researchers, see figure) (**Fig.1b**). The cultures comprised CDH1^+^ keratinocytes (**Extended Fig.1d**) without contaminating fibroblasts or immune cells (Red and white cells **in Extended Fig.1e-g**). Unlike primary keratinocytes plated on tissue culture plastic (**Extended Fig.1h**), epithelioids formed a 3D tissue-like structure with a basal layer of TRP63^+^ cells and suprabasal layers of KRT4^+^ differentiated keratinocytes (**Fig.1c-d**), recapitulating the architecture of esophageal epithelium. By using larger membranes, a single mouse esophagus could be amplified 332 x, yielding 264 cm^2^ of cultured epithelium in 30 days (**Extended Fig.1i-j**).

**Figure 1.**
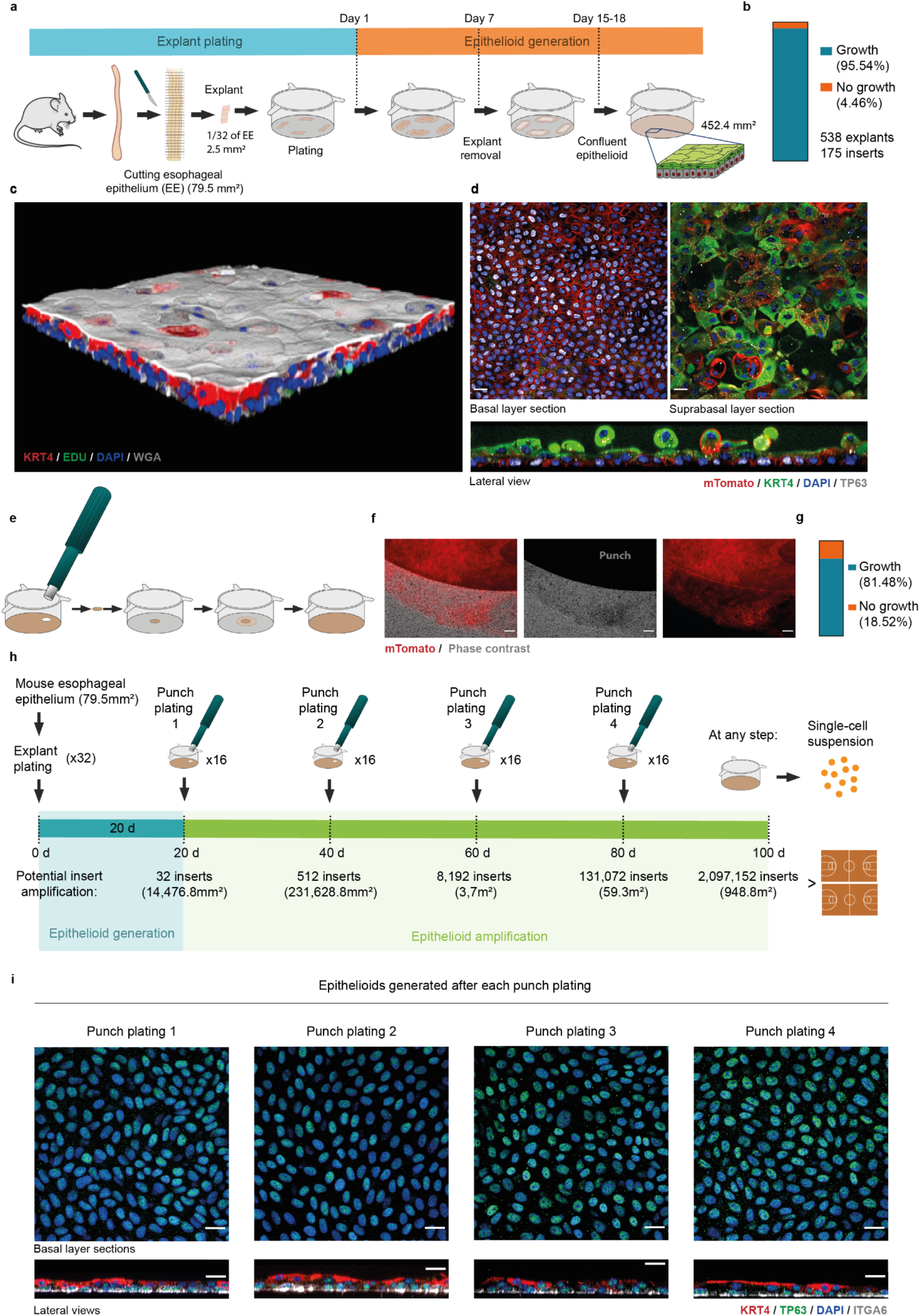
Generation and amplification of mouse esophageal epithelioids. **a**, Protocol: Epithelioid generation from four esophageal explants. **b**, Proportion of explants that contribute to epithelioid generation. *n=*538 explants from 33 mice plated in 175 inserts by 5 different researchers. **c**, Rendered confocal z-stack of a typical confluent epithelioid after 1h incubation with EdU. Stained for KRT4 (red, suprabasal cells), wheat germ agglutinin (white), EdU (green, proliferating cells) and DAPI (blue). **d**, Optical confocal sections of an mTmG mouse esophageal epithelioid showing membrane Tomato fluorescence (mTomato, red), stained for TP63 (grey), DAPI (blue) and KRT4 (green). Top left panel shows a basal cell layer section. Top right panel shows a section of suprabasal cell layers. Bottom panel shows lateral view. Scale bars, 20 μm. **e**, Scheme of the punch plating procedure. **f**, Epifluorescence microscopy images showing a front of mTmG cells exiting a punch to colonize the insert. Phase contrast (grey) and mTomato (red). Scale bar, 100 μm. **g**, Proportion of epithelioid punch biopsies able to generate new cultures. **h**, Protocol for punch plating amplification. **i**, Optical sections of basal cell layer (top panels) and lateral views (bottom panels) from confocal 3D image stacks of epithelioids stained for TRP63 (green), KRT4 (red), ITGA6 (grey) and DAPI (blue). Scale bars, 20 μm. Source data in Supplementary Table 1.

To further amplify primary epithelial cultures, the standard approach is to use enzymatic digestion to prepare a single cell suspension which is then used to establish multiple new cultures. To avoid trypsinization-induced cell damage^32^, we instead opted to expand the cultures by repeating explant process (**Fig.1e-g**). Confluent cultures were dissected into 16 pieces, each of which was placed on a permeable membrane insert. After 2-3 days cells colonized the new insert reaching confluence within 20 days (Methods, **Fig.1e-f**). The process was repeated over 4 consecutive passages and confluent stratified cultures with TRP63^+^, ITGA6^+^ progenitors overlaid by KRT4^+^ differentiated cells continued to form (**Fig.1h-i**). Using this approach, a single mouse esophagus epithelium could potentially be amplified 12×10^6^ fold in 100 days, obtaining up to 2.1×10^6^ primary cultures (of 4.5 cm^2^ each), corresponding to the area of two basketball courts (**Fig.1h**). At any stage, if required, cells can be trypsinized for downstream analysis, genetic modification and re-plating, cryopreservation or organoid generation (**Fig.1h, Extended Data Figure 1k-m**).

Long-term expansion of primary normal human esophagus cells has been challenging ^33^. Using the same method developed for mouse, we were able to generate epithelioid cultures from human esophageal epithelium of transplant donors aged 36 to 76 years (**Fig.2a-b**). Cultures maintained in cFAD expanded to confluence with high efficiency (**Fig.2c**). At confluence, the cultures had a basal layer of proliferating ITGA6^+^ keratinocytes with suprabasal layers of KRT4^+^ differentiated keratinocytes (**Fig.2d-e**). Thus, epithelioid cultures can amplify human esophageal samples from normal adult tissue and provide a robust platform for studying human esophageal biology.

**Figure 2.**
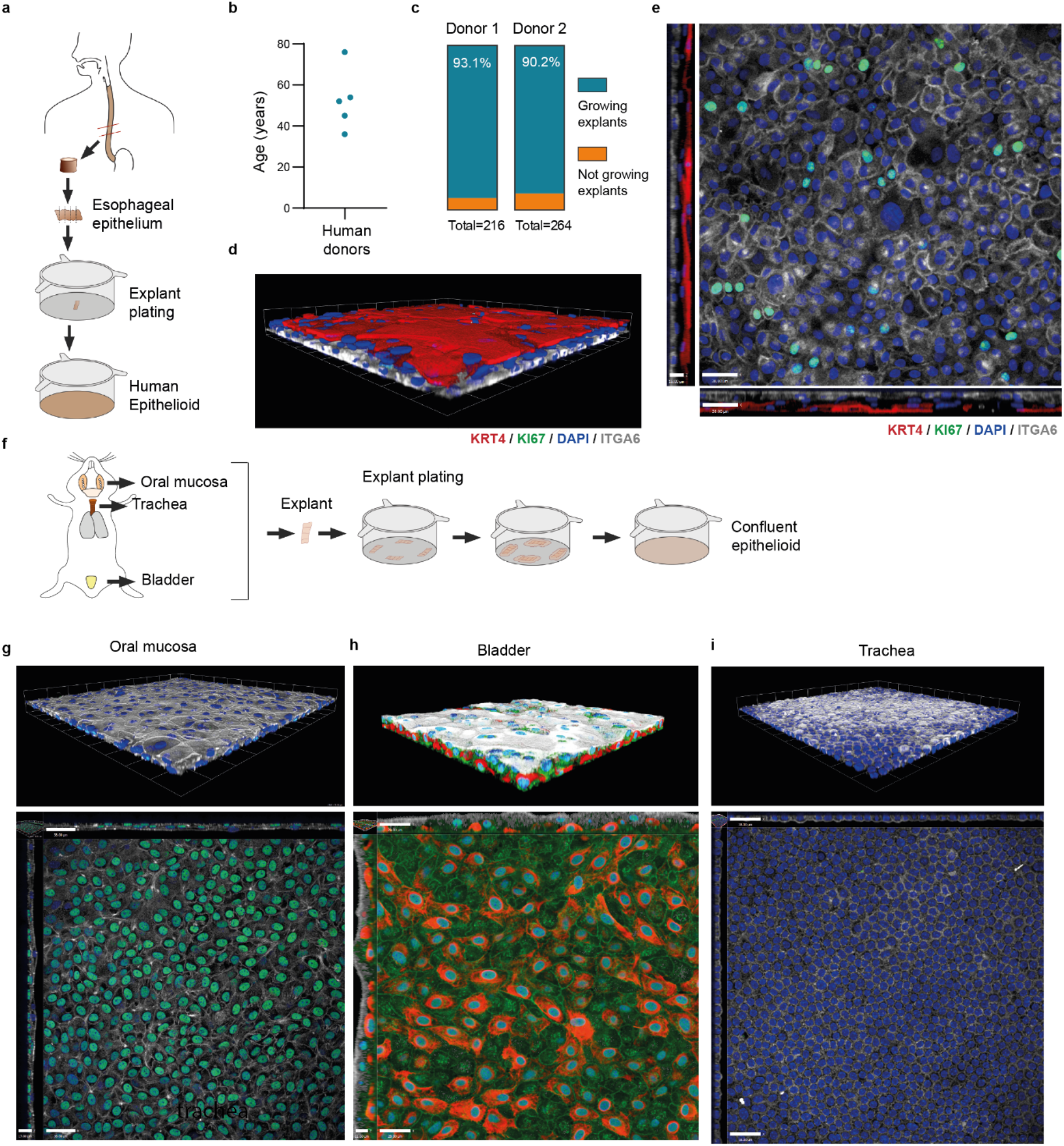
Generation of human esophageal epithelioids and mouse oral, bladder, and tracheal epithelioids. **a**, Schematic illustration of epithelioid generation from human esophagus. **b**, Age distribution of human donors expanded as epithelioids. Each dot represents a donor. **c**, Proportion of explants that form a cellular halo and contribute to epithelioid generation. Total number of explants plated per donor is indicated. *n*=480 explants from 2 donors. **d-e**, Rendered confocal z stack (d) and basal plane optical section with orthogonal views (e) of a typical human esophageal epithelioid stained for KRT4 (red), ITGA6 (white), KI67 (green) and DAPI (blue). Scale bars=38 μm for x-y and 15 μm for z. **f**, Illustration of mouse epithelioid generation from mouse oral mucosa, bladder urothelium and tracheal epithelium. **g**, Rendered confocal z stack (top) and basal layer optical section with orthogonal views (bottom) of a typical mouse oral mucosa epithelioid stained for WGA (white), TRP63 (green) and DAPI (blue). Scale bars=38 μm for x-y and 17 μm for z. **h**, Rendered confocal z stack (top) and basal plane section with orthogonal views (bottom) of a typical mouse bladder epithelioid stained for KRT20 (white), KRT5 (green), KRT14 (red) and DAPI (blue). Scale bars=28 μm for x-y and 11 μm for z. **i**, Rendered confocal z stack (top) and basal plane section with orthogonal views (bottom) of a typical mouse tracheal epithelioid stained for WGA (white) and DAPI (blue). Scale bars=38 μm. Source data in Supplementary Table 1.

We next applied the same protocol used for the esophagus to another mouse stratified squamous tissue (oral mucosa), a transitional epithelium (bladder urothelium) and a pseudostratified columnar epithelium (tracheal epithelium). In each case, the protocol generated confluent cultures that recapitulated the architecture of the original tissue (**Fig.2f-h**). These results suggest that the epithelioid system can be extended to multiple types of epithelia.

### Characterization of esophageal epithelioid cultures

We went on to characterize mouse esophageal epithelioid cultures in depth. Explants were plated on insert membranes and when a large cellular halo had formed around the explant (around 7 days after plating) they were removed. Once a confluent stratified culture was obtained (around 15-18 days after plating), we maintained the cultures in reduced growth factor media (mFAD, methods), refreshed 2-3 times a week (**Fig.3a**). Under these conditions, epithelioids maintained the morphology of the *in vivo* tissue, with a basal layer of TRP63^+^ epithelial progenitor cells. Basal cells showed the typical polarization seen *in vivo*, expressing the hemidesmosome protein ITGA6 exclusively on the part of the membrane in contact with the basement membrane. Proliferation was restricted to the basal layer while suprabasal layers expressed KRT4 cells recapitulating *in vivo* features (**Fig.3b-e**). The proportion of basal cells in S-phase was 11%, comparable to that of the adult mouse esophagus (**Extended Fig.2a-c**). Cell tracing of proliferating cells and confocal live imaging showed that in epithelioids, similarly to the *in vivo* tissue, basal cells proliferate, and some exit the basal layer migrate into the suprabasal layers and are eventually shed (**Extended Fig.2d-f, Supplementary videos 2-4 and Fig.3f-i**).

**Figure 3.**
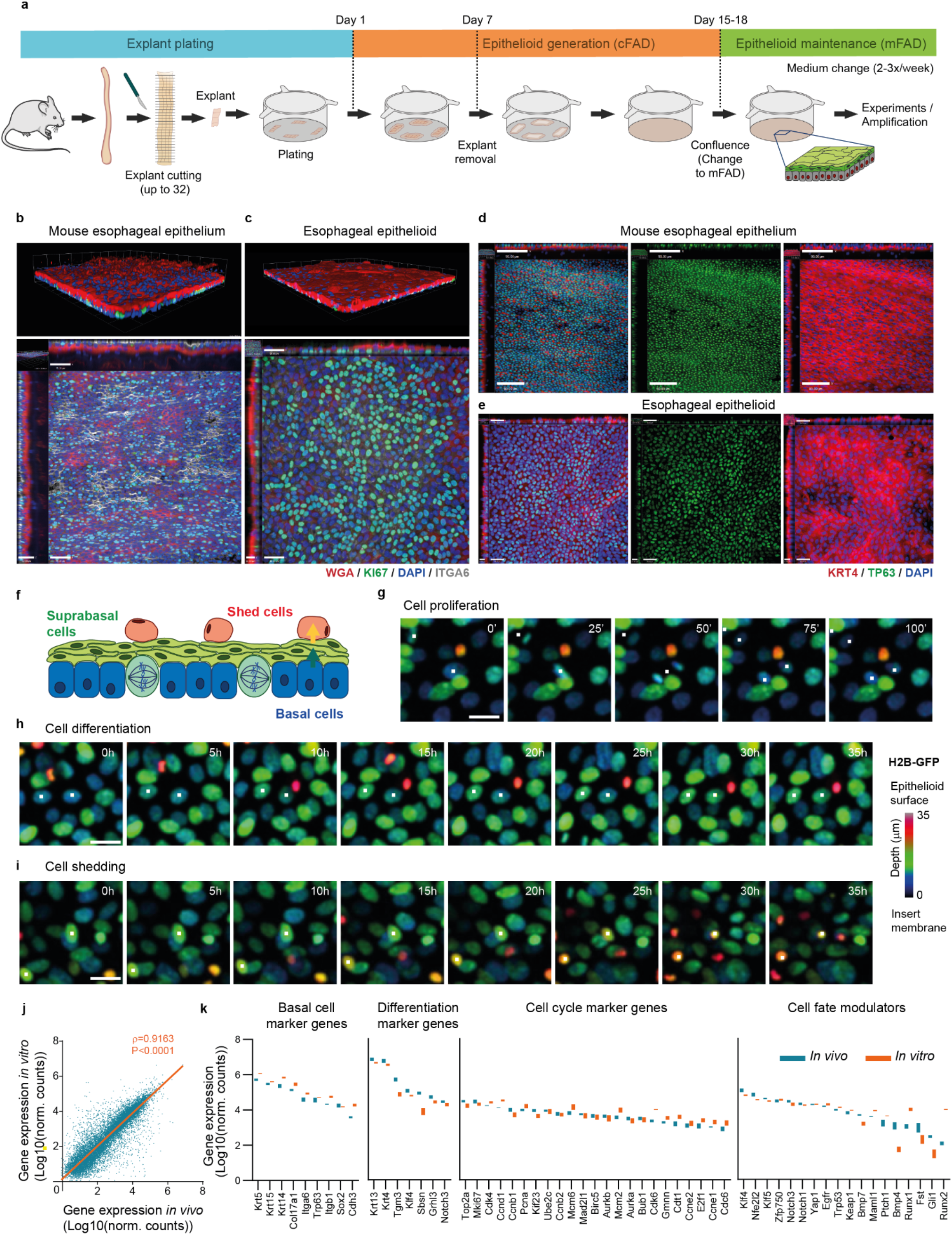
Characterization of mouse esophageal epithelioids. **a**, Schematic illustration of epithelioid generation from four esophageal explants and its maintenance at confluence. **b-c**, Rendered confocal z stack (top) and basal layer optical section with orthogonal views (bottom) of a typical mouse esophagus wholemount (b, scale bars=41 μm for x-y and 32 μm for z) and esophageal epithelioid (c, scale bars=38 μm for x-y and 16 μm for z) stained for ITGA6 (white), KI67 (green), KRT4 (red) and DAPI (blue). **d-e**, Basal layer optical section with orthogonal views (bottom) of a typical mouse esophagus wholemount (d, scale bars=40 μm for x-y and 24 μm for z) and esophageal epithelioid (e, scale bars=38 μm for x-y and 15 μm for z) stained for TP63 (green), KRT4 (red) and DAPI (blue). **f-i**, Confocal live imaging of H2BGFP expressing epithelioids showing multiple z-projection time frames labelled with a z-depth rainbow colour scale, where colour indicates cell position in z-plane. (**f)** Scheme of esophageal epithelioid structure with the z-scale colour labelling used in **g-I**, with basal cells (blue), suprabasal cells (green) and shedding cells (red). Selected live images showing cells undergoing mitosis (**g**), differentiation (**h**) and shedding (**i**) from Supplementary videos 2-4 respectively. Time is indicated in each frame. Scale bars, 20 μm. **j-k**, RNA sequencing comparing gene expression from mouse esophageal epithelium (*in vivo*) and esophageal epithelioids (*in vitro*). *n=*4 animals and 4 epithelioids from 4 different animals. Correlation of all transcripts (**j**) and selected basal cell markers, differentiation marker genes, cell cycle marker genes and cell fate modulators (**k**)^34^. The orange line shows the linear regression between samples with the Pearson’s coefficient and p-value of the correlation. Source data in Supplementary Table 1.

RNA-sequencing analysis showed that gene expression in epithelioids highly correlated with the gene expression of epithelial cells directly isolated from mouse esophagus (**Fig.3j**). The expression of epithelial transcripts (*Epcam, E-cadherin, Zo-1*), (**Extended Fig.2g**), mRNAs characteristic of basal, differentiating and cycling cells, and of multiple esophageal cell fate regulators was not substantially altered by the culture conditions (**Fig.3k**)^29,34-40^, arguing that *in vivo* cell fate regulation may be preserved in epithelioids. Analysis of gene signatures of keratinocyte differentiation states^34^ showed a correlation between *in vivo* and *in vitro* with the largest differences confined to late differentiation markers, reflecting decreased terminal differentiation in epithelioids (**Extended Fig.2h**). Air-liquid interface culture^16,41^, which enhances terminal differentiation, may be used to correct this discrepancy (**Extended Fig.2i-k**).

### Long-term epithelioids maintain tissue turnover and remain genomically stable

To model *in vivo* esophageal tissue, epithelioid cultures should be able to self-maintain in the long-term without passaging. We found epithelioids kept in mFAD refreshed twice a week remained in a steady-state for one year, with approximately constant levels of cell density and cell proliferation (**Fig.4a-d**). As epithelial cells may also develop copy number alterations when expanded *ex vivo* (CNA) ^43^, we performed whole-genome sequencing, finding only a subpopulation of cells (17-29%) with detectable CNA after 8 months in continuous culture (**Fig.4e-f and Extended Fig.3a-c**).

**Figure 4:**
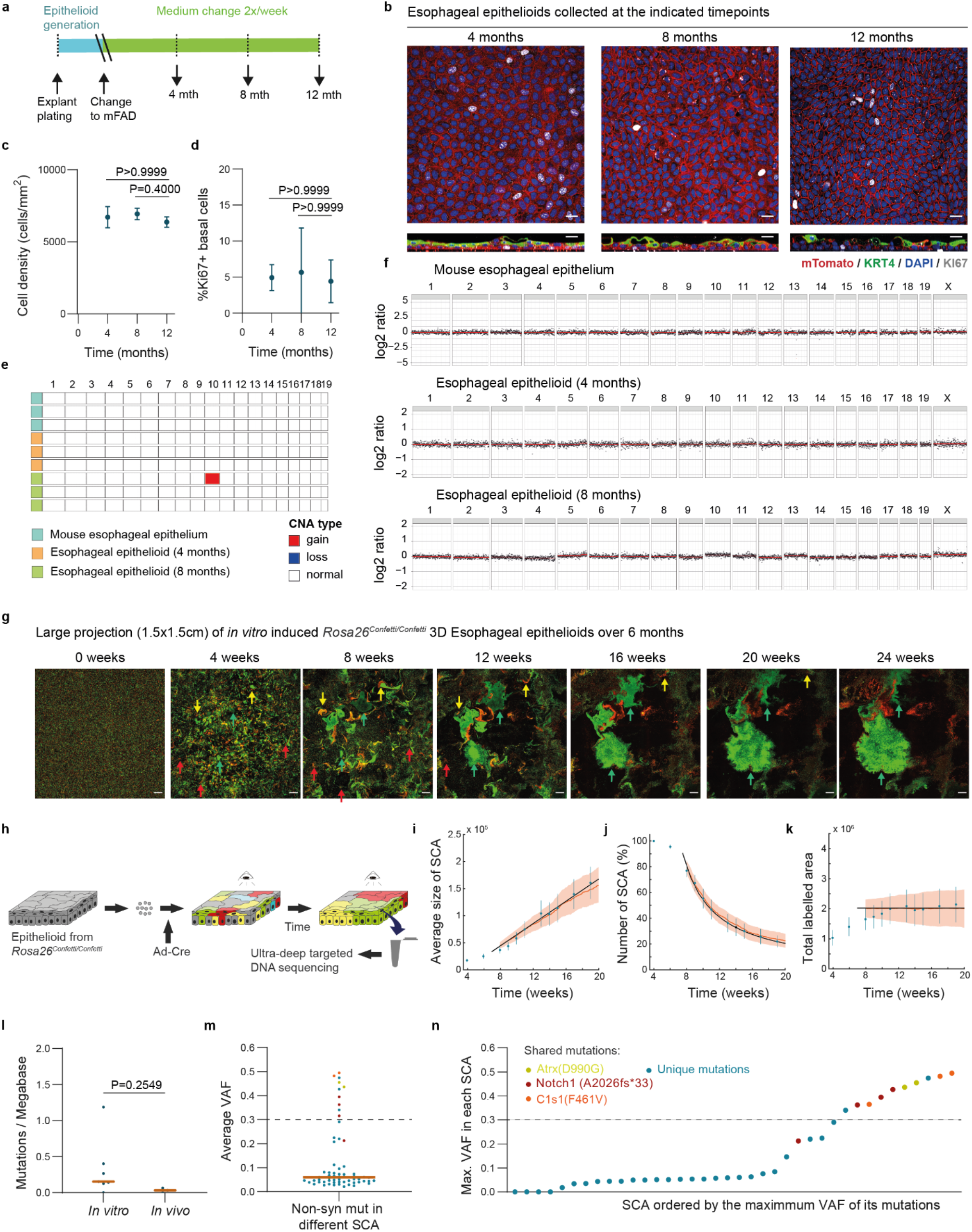
Long-term maintenance and tissue dynamics of esophageal epithelioids. **a-d**, Esophageal epithelioids generated from *Rosa26*^*mTmG*^ mice maintained without passaging at different for up to 12 months and stained for KI67 (grey, proliferating cells), KRT4 (green, differentiated cells) and DAPI (blue). **a** protocol and **b** optical confocal section of the basal cell layer (top panels), lateral views (bottom panels). Scale bars, 20 μm. **c-d**, Cell density (**c**) and the proportion of KI67^+^ basal cells (**d**). Mean±S.D are shown, *n*=3 inserts from different animals per time point. Two-tailed Mann-Whitney test. **e-f**, Whole genome sequencing of mouse esophageal epithelium and epithelioids after 4 and 8 months in culture. *n=*3 animals and 3 esophageal epithelioids from different animals per time point. Summary plot showing all gain and loss of chromosome regions that affect more than 20% of cells (**e**) and copy number profile (**f**). **g-k**, Epithelioids from Rosa26^confetti/wt^ mice cultured for 24 weeks after *in vitro Cre* recombination (see methods). **g**, Representative images of the same region of an epithelioid at the indicated time points. Scale bars= 1mm. Red arrows indicated shrinking single-colored areas (SCA), yellow arrows indicate SCA with biphasic growth and green arrows indicate growing SCA. **h**, Experimental protocol for **g** and clone cutting and DNA-sequencing used in **l-n. i-k**, Average SCA size (**i**), SCA number (**j**) and total labelled area (**k**) with experimental values (in blue, mean^+^-SD) and a theoretical, single-parameter fit (in black) as well as lattice-based simulations (in orange) of a single-progenitor model. *n=* 351 SCA from 9 epithelioids from 6 different animals. **l-n**, 38 surviving SCA were collected by laser-capture microscope and DNA was sequenced. Estimated mutation burden of the SCA collected (*in vitro*) and 3 control mice samples (*in vivo*) (l), average VAF of non-synonymous mutations in different SCA (**m**) and SCA ordered by the maximum VAF of its mutations, mutations represented in more than one sample are highlighted in the specified colors are shown (**n**). Orange bars indicate mean values, and dashed lines a VAF threshold for clonal mutations in the sample. Unpaired two-sided Student’s *t*-test. Source data in Supplementary Table 1.

### Long-term cell dynamics in epithelioids

We next analyzed long-term cell behavior in epithelioids by lineage tracing. Epithelioids were generated from *R26-confetti* mice, multicolor heritable cell labelling induced with adenoviral *Cre* recombinase, and the labelled cells placed in epithelioid culture for 6 months without passaging (methods). The cultures were imaged weekly using an Incucyte live imaging system (Essen Bioscience) (**Fig.4g-h**). Initially, the culture is formed by a mixture of differentially colored individual cells. However, after 1 month, single-colored areas (SCA) started appearing, which are generated from a single cell or neighboring cells with the same color. We followed the behavior of 351 SCA from 9 cultures coming from 6 different animals starting 5 weeks after plating. SCA of sufficient size to be resolved at 5 weeks were tracked. We observed different patterns of evolution of SCA area. Most SCA became smaller (80 %), others grew and then shrunk (12 %), a minority remained constant in area (1 %) and finally some SCA grew progressively (7 %) (**Extended Fig.3d-f**). Furthermore, from 8 weeks, the number of SCA declined, but the average size of the remaining SCA increased, so that the total labelled area remained approximately constant (**Fig.4i-k**). These features are hallmarks of neutral drift, observed in clones labelled with a neutral reporter in squamous epithelia *in vivo* ^11,44^. Two simple quantitative models of cell behavior in esophageal epithelium gave a good fit to the data (Methods, **Fig.4i-k**)^45^. As the behavior of SCA recapitulates that of neutral clones *in vivo* we conclude progenitor dynamics in epithelioids is similar to those in mouse esophagus *in vivo* ^11,45^.

As animals age, they accumulate somatic mutations that may result in clonal expansions if they affect genes that regulate progenitor cell fate^46^. This process occurs at a low rate in the esophagus of ageing wild type mice, where mutations in genes such as *Notch1* are seen in expanded clones in mice of 1 year of age ^19,25^. To determine if somatic mutations impact the growth of SCA, 46 samples from 38 surviving SCA at the 9 month time-point were isolated by laser-capture microdissection. DNA was extracted and targeted sequencing performed for 192 genes implicated in driving clonal expansions in squamous epithelia and/or recurrently mutated in squamous cancer^19^. Median coverage was 106 fold. The estimated mutational burden was similar to that in age matched mouse oesophagus^19^, arguing the mutation rate is not substantially increased in epithelioid culture (**Fig.4l**). The low variant allele frequency (VAF) of most mutations, (71 % mutations had VAF<0.1, **Fig.4m**), indicates that these were unlikely to have altered SCA dynamics, as to do so a mutation must have a VAF close to 0.5 (at which level most cells in the SCA carry the mutation assuming they are diploid). This was the case for 26 % of SCA (**Fig.4n**), with 80 % of these sharing mutations with other SCA, suggesting that most mutations with large VAF were already present before labelling. Interestingly, the mutation shared by more SCA was a *Notch1* frameshift mutation. This is consistent with the development of spontaneous *Notch1* mutations that drive clonal expansions in ageing mice^19,25^. Therefore, most of SCA behavior can be explained by neutral drift and are not caused by the acquisition of a driver mutations *in vitro*.

### Epithelioids as a tool to study neutral and non-neutral cell competition in stratified epithelia

The properties of epithelioids led us to speculate that they may be suitable for studying slow processes such as clonal competition in a tissue-like environment. We first investigated neutral competition between two populations of equal fitness. We established esophageal epithelioids from conditional *R26-EYFP* mice in which cells and their descendants express Enhanced Yellow Fluorescent Protein (EYFP) after genetic recombination by *Cre* recombinase^11^. Cells were infected with adenovirus encoding *Cre* achieving a 90±1 % recombination rate (**Extended Fig.4a**). RNAseq showed that the only transcript significantly altered by recombination was *Rosa26* mRNA (5.17 fold change, adjusted p-value 1.52×10^−72^) (**Extended Fig.4b**). Thus Cre-mediated *loxP* excision can be performed at high efficiency without altering overall gene expression. We then generated mixed epithelioid cultures with EYFP^+^ recombined and unrecombined cells from the same esophagus and measured the proportion of each subpopulation over time (**Extended Fig.4c and Fig.5a-b**). The proportion of EYFP^+^ cells remained constant over 2 months (**Fig.5c**). This recapitulates the neutral behavior of the same reporter allele in the esophagus *in vivo*^11^.

**Figure 5:**
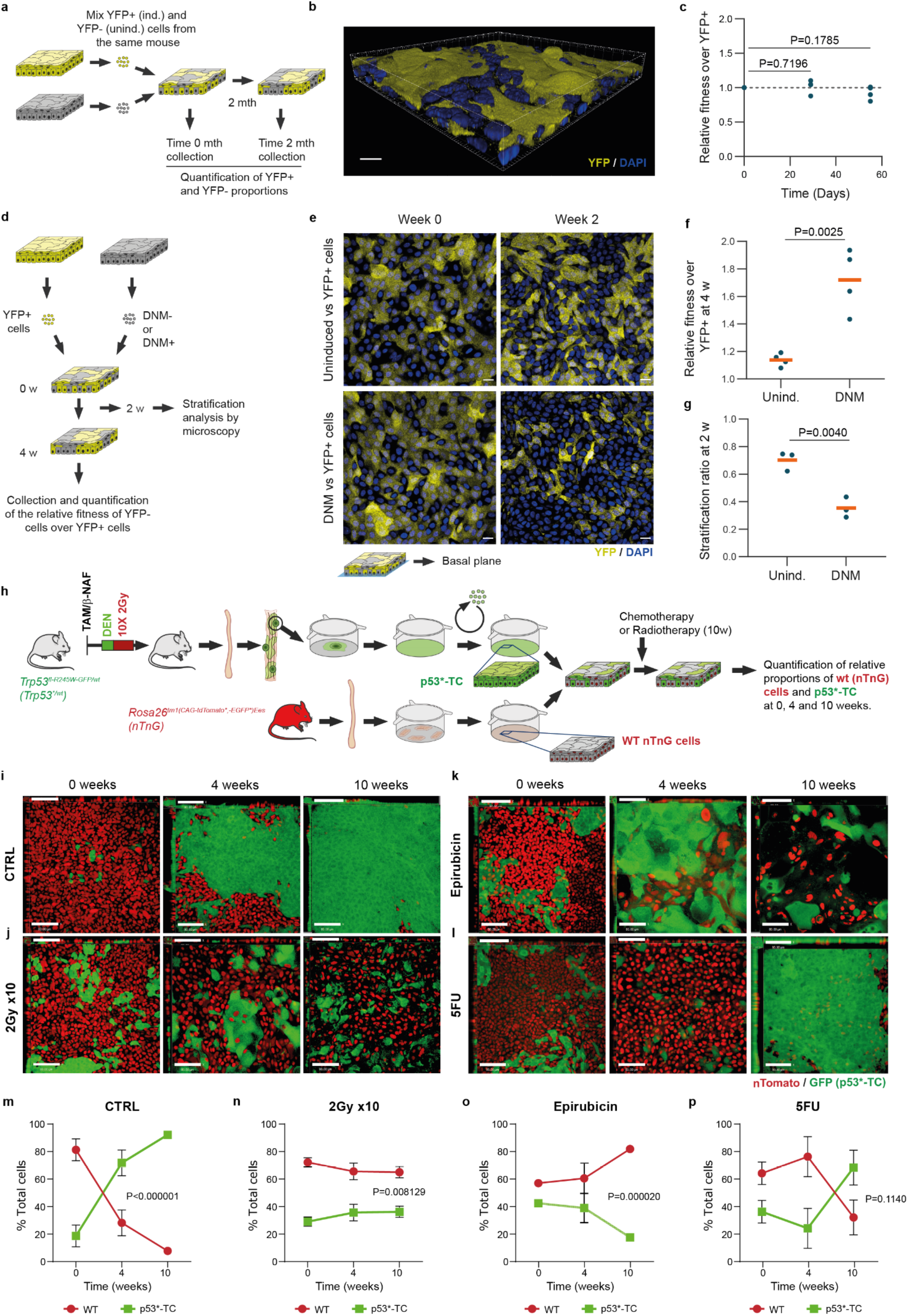
Epithelioids as a tool to study clonal competition. **a-c**, Cell competition in mixed epithelioid cultures formed by induced (YFP^+^) and uninduced (YFP-) *Rosa26*^*YFP/YFP*^ cells were maintained for 2 months. **a**, Protocol scheme. **b**, Rendered confocal z stack of a typical epithelioid with both competing subpopulations. **c**, Relative fitness of YFP-versus YFP^+^ cells at different time points. *n=*4 inserts per time point from 4 different animals. Unpaired two-sided Student’s t-test. **d**, Protocol for Dominant Negative Mastermind like 1 (DNM) and wild type cell competition. Primary cells from *R26*^*flDNM*^ mice epithelioids, uninduced (wild type, DNM-) or induced (DNM^+^) *in vitro*, were mixed with YFP^+^ cells and kept for in culture. **e**, optical sections of basal cell plane at 0 and 2 weeks of competition of conditions shown in d, Scale bar, 20 μm. **f-g**, Relative fitness over YFP^+^ cells (f, *n=*4) at 4 weeks and stratification ratio of DNM- and DNM^+^ cells at 2 weeks (g, *n=*3). Replicates correspond to primary cultures from different animals. Orange lines show mean values. Unpaired two-sided Student’s t-test. **h-p**, Co-culture of transformed and wild type cells. **h**, Protocol to generate p53 mutant transformed cells (p53*-TC). *p53*^*R245W*^ mutant esophageal tumors were generated and expanded using the epithelioid protocol (see methods) to obtain p53*-TC. p53*-TC cells were mixed with primary wild type cells from *Rosa26*^*nTnG*^ mice and exposed to 10 weeks of weekly dosing with 2 Gy gamma irradiation, 1µM epirubicin or 5µM 5-fluorouracyl (5FU). **i-l**, Orthogonal views of basal layer optical sections of z-stacks at 0, 4 and 10 weeks of treatment. Scale bars, 80 μm. **m-p**, Proportion of each subpopulation over time. *n=*3. Mean±SEM are indicated. P-values indicate comparison between subpopulations at the 10-week time point. Unpaired two-sided Student’s t-test. Source data in Supplementary Table 1.

Next, we studied a non-neutral competition. We selected a conditional dominant negative mutant of *Maml1* (DNM) that has strong advantage over wild type cells in the esophageal epithelium *in vivo*^47^. Epithelioids were generated from *R26-DNM* mice and infected with either null or *Cre*-encoding adenovirus to generate wild-type or DNM expressing keratinocytes from the same mice. These cells were mixed with EYFP expressing cells as described above and the proportions of cells were analyzed after 4 weeks (**Fig.5d**). DNM expressing cells outcompeted EYFP^+^ cells, showing a significant increase in relative fitness relative to the uninduced cells from the DNM mice (**Fig.5e-f**). Confocal microscopy of day 15 cultures showed that ratio of suprabasal: basal cells was significantly lower for DNM expressing than non-expressing cells (**Fig.5g**). This is consistent with the behavior of DNM expressing clones *in vivo*, which gain a competitive advantage by progenitors generating fewer differentiating than progenitor daughters per average cell division ^47^. We conclude that epithelioids are suitable platform for studying mutant cell competition.

### Effect of chemotherapy and radiotherapy on the competition between transformed and non-transformed cells

Next, we turned to a more challenging application. Many cytotoxic cancer treatments cause substantial normal tissue damage alongside tumor cell killing. We hypothesized that longevity of epithelioids may allow mixed transformed/normal cell co-cultures to undergo a prolonged course of treatment replicating studies currently performed *in vivo*. In such experiments the normal and transformed cells compete in the same well with the net benefit/detriment of treatment emerging from the proportion of surviving cells of each type.

We began by generating transformed cells (p53*-TC), collecting, and expanding cells from a tumor generated in a mutant mouse carrying an inducible *Trp53*^*R245W*^ allele (see methods), equivalent to the human cancer hotspot *TP53*^*R248W*^ mutation. These cells carry substantial CNA including translocations, telomeric associations, duplications and trisomies of individual chromosomes and 55 % of cells were near tetraploid (**Extended Fig.4d-e**). The transformed cells were mixed with wild type cells from a *Rosa26*^*nTnG*^ mouse which express the Tomato protein targeted to the cell nucleus in epithelioid cultures. The transformed cells outcompeted the wild type cells (**Fig.5i and m**).

Next, we investigated the impact of anti-cancer treatments on wild type and transformed cell competition by exposing the cultures to intermittent doses of ionizing radiation (IR), epirubicin or 5-fluorouracyl (5FU) (**Fig.5j-p and Extended Fig.4e**), which are used to treat esophageal cancer ^48^. All three treatments altered the competition between wild type and transformed cells over 10 weeks. 2 Gy IR exposure halted the expansion of transformed cells and induced aberrant large nuclei specifically in the transformed population (**Fig.5j and n and Extended Fig.4e**). Conversely, epirubicin showed significant toxicity in both wild type and transformed cells, though the effect on cell fitness was more pronounced in transformed cells which were progressively depleted from the culture (**Fig.5k and o and Extended Fig.4e**). 5FU treatment initially inhibited the expansion of transformed cells, however, later transformed cells recovered and overtook wild type cells (**Fig.5l and p and Extended Fig.4e**), consistent with the development of 5FU resistance published using other *in vitro* models^49^.

These results show the potential of epithelioids to study the differential effects of therapy on transformed and wild type cells competing in long-term continuous co-culture.

## Discussion

The epithelioid system emerges as a facile and versatile method of generating 3D sheets of cultured epithelial tissue with multiple applications. This technique allows the production and long-term maintenance of large amounts of primary 3D epithelium from a small initial sample. It may be applied to human epithelia, allowing the amplification of small, precious patient tissue biopsies for the study genetic or other disorders in an organotypic context. Murine epithelioids can be generated from genetically manipulated mice, enabling a wide range of transgenic tools and sensors to be leveraged. Epithelioids are also amenable for live imaging, facilitated by its growth on a flat surface rather than as spheroids in suspension. Genetically manipulated cells may be followed by lineage tracing, paralleling *in vivo* studies but with substantial savings in time and cost. In combination of these properties enhance the analysis of multiple processes such as progenitor cell dynamics, epithelial differentiation, or cell-cell interactions in an organotypic context.

A particular advantage of epithelioids over other advanced cell culture methods is their ability to self-sustain for weeks to months without passaging, allowing processes that evolve slowly within tissues to be studied. These include competition between mutant or transformed cells versus wild type cells, permitting experimental validation of the selection of somatic mutations, and to define the effect of drug treatments on such competition, as shown above.

More broadly, potential applications of this system extend to studying how mutagenesis^19,21^, environmental exposures such as ionizing radiation^28^, physical damage such as wounding, ageing, long-term drug treatment, inflammation or metabolic alterations affect cellular states, and tissue function in a 3D epithelial context.

## Acknowledgements

This work was supported by grants from the Wellcome Trust to the Wellcome Sanger Institute (098051 and 296194) and Cancer Research UK Programme Grants to P.H.J. (C609/A17257 and C609/A27326). A.H. benefited from the award of an EMBO long-term fellowship (EMBO ALTF885-2015) and a Maria Zambrano Grant to attract international talent from Universitat de Barcelona and Ministerio de Universidades and cofunded with Next Generation EU funds. D.F-A work was supported by funding from the European Union FP7-Euratom-Fission award 323267, Risk, Stem Cells and Tissue Kinetics – Ionising Radiation at the Wellcome Sanger Institute. M.P.A. acknowledges funding from the Wellcome Trust and The Royal Society (105942/Z/14/Z and 105942/Z/14/A), and the UKRI Medical Research Council (MR/P019013/1). J.A.V-L and I.S.F work was supported by joint National Centre 3Rs-Cancer Research UK award NC/X000885/1 and Cancer Research UK RadNet grant C17918/A28870 to D.F-A at the Gurdon Institute. G.P. is supported by the Agencia Estatal de Investigación of Spain (Grant No. PID2020-116163GA-I00). S.C.D. benefited from the award of an ESPOD fellowship, 2018-21, from the Wellcome Sanger Institute and the European Bioinformatics Institute EMBL-EBI.

We thank Esther Choolun, Tom Metcalf and Sanger RSF facility for technical support with animal research. We thank Claire Hardy, Calli Latimer, staff from the Cancer, Ageing, and Somatic Mutations programme support laboratory for technical support with sequencing and The Gurdon Institute core imaging facility for technical support with confocal microscopes.

## Author contributions

A.H., D.F.A and P.H.J designed experiments. M.P.A., performed the initial experiments and set up the 3D *in vitro* model. A.H., D.F.A., A.K., U.B., K.M., E.A., J.A.V-L and I.S.F performed experiments. G.P. and A.R.L., analyzed experimental data and performed mathematical modelling. S.C.D, C.K., and S.H.O analyzed sequencing data. K.S-P. and K.M. provided human samples. A.H., D.F.A., M.P.A. and P.H.J wrote the paper. M.G supervised the research of S.C.D and P.H.J supervised all the research. All authors discussed the results and commented on the manuscript.

## Supplementary Information

<u>Extended Figures 1-5</u>. (Embedded in this manuscript)

<u>Supplementary Table 1</u>. Source Data.

<u>Supplementary Video 1.</u>

Example of epithelioid generation from a single esophageal explant from an *Rosa26*^*mTmG*^ mouse tracked by live imaging using an ‘Incucyte’ system. mTomato fluorescence intensity is indicated using the Rainbow LUT of Fiji ImageJ. Scale bar, 1000 μm. See also **Extended Data Fig.1b**.

<u>Supplementary Video 2.</u>

Example of cells undergoing mitosis in the basal layer of H2BGFP expressing esophageal epithelioids (see Methods) imaged by confocal live imaging. The z-projection of a full 3D stack is shown using a z-depth rainbow color scale (see Fig.3), in which cells are labelled according to its z-plane position in the epithelioid. As a result, blue, green and red cells corresponding to basal cells, suprabasal cells and shedding cells, respectively. Time is indicated in each frame. Scale bars, 20 μm. Related to **Fig.3g**.

<u>Supplementary Video 3.</u>

Example of cells migrating upwards to become differentiated cells. Images taken from H2BGFP expressing esophageal epithelioids (see Methods) imaged by confocal live imaging. The z-projection of a full 3D stack is shown using a z-depth rainbow color scale (see Fig.3), in which cells are labelled according to its z-plane position in the epithelioid. As a result, blue, green and red cells corresponding to basal cells, suprabasal cells and shedding cells, respectively. Time is indicated in each frame. Scale bars, 20 μm. Related to **Fig.3h**.

<u>Supplementary Video 4.</u>

Example of cells shedding from H2BGFP expressing esophageal epithelioids (see Methods) imaged by confocal live imaging. The z-projection of a full 3D stack is shown using a z-depth rainbow color scale (see Fig.3), in which cells are labelled according to its z-plane position in the epithelioid. As a result, blue, green and red cells corresponding to basal cells, suprabasal cells and shedding cells, respectively. Time is indicated in each frame. Scale bars, 20 μm. Related to **Fig.3i**.

<u>Supplementary Video 5.</u>

Example of Single Color Area (SCA) dynamics. Epithelioids from Rosa26^confetti/wt^ mice cultured for 24 weeks after *in vitro Cre* recombination (see methods). Repeated imaging of the same region at the indicated time points using an Incucyte system. Scale bars= 1mm. Related to **Fig.4g**.

<u>Supplementary Video 6.</u>

Example of an expanding Single Color Area (SCA). Epithelioids from Rosa26^confetti/wt^ mice cultured for 24 weeks after *in vitro Cre* recombination (see methods). Repeated imaging of the same region at the indicated time points using an Incucyte system. Scale bars= 1mm. Related to **Extended Data Fig.3d**.

<u>Supplementary Video 7.</u>

Example of a biphasic Single Color Area (SCA). Epithelioids from Rosa26^confetti/wt^ mice cultured for 24 weeks after *in vitro Cre* recombination (see methods). Repeated imaging of the same region at the indicated time points using an Incucyte system. Scale bars= 1mm. Related to **Extended Data Fig.3d**.

<u>Supplementary Video 8.</u>

Example of a biphasic Single Color Area (SCA). Epithelioids from Rosa26^confetti/wt^ mice cultured for 24 weeks after *in vitro Cre* recombination (see methods). Repeated imaging of the same region at the indicated time points using an Incucyte system. Scale bars= 1mm. Related to **Extended Data Fig.3d**.

<u>Supplementary Video 9.</u>

Example of a shrinking Single Color Area (SCA). Epithelioids from Rosa26^confetti/wt^ mice cultured for 24 weeks after *in vitro Cre* recombination (see methods). Repeated imaging of the same region at the indicated time points using an Incucyte system. Scale bars= 1mm. Related to **Extended Data Fig.3d**.

<u>Supplementary Video 10.</u>

Example of a shrinking Single Color Area (SCA). Epithelioids from Rosa26^confetti/wt^ mice cultured for 24 weeks after *in vitro Cre* recombination (see methods). Repeated imaging of the same region at the indicated time points using an Incucyte system. Scale bars= 1mm. Related to **Extended Data Fig.3d**.

## METHODS

### Animals

Multiple strains were used as a tissue source. C57/Bl6 mice were used as wild type, unless specified. In addition, we used the following genetically engineered mouse strains from the Jackson Laboratory: *Rosa26*^*mT/mG*^ (RRID:IMSR_JAX:007676)^50^, *Rosa26*^*M2rtTA*^*/TetO-H2BGFP* mice^51^, doubly transgenic for a reverse tetracycline-controlled transactivator (rtTA-M2) targeted to the *Rosa26* locus and a HIST1H2BJ/EGFP fusion protein (H2BGFP) expressed from a tetracycline promoter element (RRID:IMSR_JAX:005104)^11,52^, multicolor reporter line *Rosa26*^*tm1(CAG-Brainbow2*.*1)Cle*^ (*R26-confetti*, RRID:IMSR_JAX:017492)^53^ *Rosa26*^*flYFP/flYFP*^ mice (R26-YFP, RRID:IMSR_JAX:006148)^54^, *Rosa26*^*nT/nG*^ (RRID:IMSR_JAX:023035), *Nfe2l2*^*tm1Ywk*^ (RRID:IMSR_JAX:017009), *Notch1*^*fl/fl*^ (RRID:IMSR_JAX:007181)^55^, *LSL Kras*^+/*G12D*^ (RRID:IMSR_JAX:019104)^50^ and *Rosa26*^*flDNM-GFP/wt*^ (RRID:IMSR_JAX:032613*R26-DNM*)^47,56^. The other mouse strains used were *Trp53*^*flR245W-GFP/wt*^ (European Mutant Mouse Archive, EM:13118)^30^ and *Ahcre*^*ERT* 57^. The mouse experiments ethics was reviewed and approved by the Welcome Sanger Institute Ethics Committee and experiments conducted according to UK government Home Office project licences PPL22/2282, PPL70/7543, and PPL4639B40. Both male and female mice between 10-16 weeks of age at the start of the experiments were used. Animals were housed in individually ventilated cages and fed on standard chow. Mice were maintained at SPOF health status.

### Epithelioid generation and maintenance

Mice were euthanized, esophagus, trachea, bladder or oral mucosa were collected and the muscle layer removed with forceps, epithelium was cut in pieces (up to 32 for a mouse esophagus) and placed on top of a transparent ThinCert™ insert (Greiner Bio-One, 657641) with the epithelium facing upward and the submucosa facing the membrane. Where indicated explants were plated on top of 44cm2 inserts (Corning, 3419). Complete FAD (cFAD) was used to culture the cells until the epithelioids reach confluence. cFAD contained DMEM (Invitrogen, 11971-025):DMEM/F12 (Invitrogen, 31330-038) (1:1), supplemented with 5 % fetal calf serum (PAA Laboratories, A15-041), 5% Penicillin-Streptomycin (Sigma Aldrich, P0781), 5 μg/ml insulin (Sigma-Aldrich I5500), 1.8×10^−4^ M adenine (Sigma-Aldrich, A3159), 1×10^−10^ M cholera toxin (Sigma-Aldrich, C8052), 10 ng/ml Epidermal Growth Factor (EGF, PeproTech EC, Ltd 100-15), 0.5 μg/ml hydrocortisone (Calbiochem, 386698) and 5 μg/ml Apo-Transferrin (Sigma-Aldrich, T2036). 7 days after plating, explants were removed by aspiration. Media was changed every three/four days. Once confluent, mouse esophageal epithelioids are maintained in minimal FAD (mFAD), containing DMEM (Invitrogen, 11971-025):DMEM/F12 (Invitrogen 31330-038) (1:1), supplemented with 5 % fetal calf serum (PAA Laboratories, A15-041), 5 % Penicillin-Streptomycin (Sigma Aldrich, P0781), 5 μg/ml insulin (Sigma-Aldrich, I5500) and 5 μg/ml Apo-Transferrin (Sigma-Aldrich, T2036). Where indicated epithelioids were lifted to the air-liquid interface by removing the media on top of the insert and maintained for 15 days.

### Immunofluorescence

For whole-mount staining, the mouse esophagus was opened longitudinally, the muscle layer removed and the epithelium incubated for 3h in 5 mM EDTA-PBS at 37°C. The epithelium was peeled from submucosa and fixed in 4% paraformaldehyde in PBS for 30 min. For epithelioid staining, inserts were washed with PBS and fixed in 4% paraformaldehyde in PBS for 30 min. Then, tissue whole-mounts or membrane inserts were blocked for 1 h in blocking buffer (0.5% bovine serum albumin, 0.25% fish skin gelatin, 1% Triton X-100 and 10% donkey serum) in PHEM buffer (60 mM PIPES, 25 mM HEPES, 10 mM EGTA, and 4 mM MgSO_4_·7H_2_O). All reagents were purchased from Sigma Aldrich. Tissues were incubated with primary antibodies (**Supplementary table 1**) overnight using blocking buffer, followed by 4 washes with 0.2 % Tween-20 in PHEM buffer of a minimum 15 min. When indicated EdU incorporation was detected with Click-iT chemistry kit following the manufacturer’s instructions (Life technologies, 23227). Next, whole-mounts or inserts were incubated overnight with 1 μg/ml DAPI (Sigma Aldrich, D9542) and secondary antibodies (1:500) in blocking buffer. When indicated Alexa fluor 647-wheat germ agglutinin (WGA, Invitrogen W32466) was added 1:200 and Alexa fluor 647 anti-human/mouse CD49f (Biolegend, 313610) was added 1:250. Afterwards, samples were washed 4×15 min with 0.2% Tween-20 in PHEM buffer and mounted in Vectashield (Vector Laboratories, H-1000). Imaging was performed using a SP8 Leica confocal microscope with a 40 x objective with 1x digital zoom, optimal pinhole and line average, bidirectional scan, speed 400-600 hz, resolution 1024×1024. 3D stacks were generated including all the cell layers of the culture and where indicated, the basal layer plane was selected. Rendered confocal z-stacks were generated using Volocity 6.3 (Perkin Elmer) and Imaris 4.3 (Bitplane). Orthogonal views and individual planes were generated using Volocity 6.3 (Perkin Elmer) or Fiji ImageJ ^58^.

### Live imaging

IncuCyte live-cell imaging system (Essen Bioscience) was used for whole-well imaging once a day (mTmG growth curve experiments) or once a week (confetti lineage tracing experiments) using its 4 x objective. Images were analyzed using the Fiji Image J software ^58^.

To evaluate cell division and cell differentiation, cells from *Rosa26*^*M2rtTA*^*/TetO-H2BGFP* mice^59^ were cultured in 6-well inserts. Confluent epithelioids were treated with doxycycline for 5 days to induce H2BGFP expression. Inserts were placed into custom-made holders to adapt them to a Leica SP8 confocal microscope stage. Cells were imaged using a HC PL FLUOTAR 40x/0.6 dry objective taking images of 512×512 resolution at 700 hz using a 1.28 x zoom, 0.5 AU pinhole, and a line average of 2. 32-plane z-stacks were obtained every 25 min for up to 40 h. Timelapses were analyzed using Fiji ImageJ software.

### Organoid generation

To generate organoids, wild-type or C57BL/6 mouse esophageal epithelioids were washed in PBS and incubated with 0.05 % Trypsin-EDTA for 20 min at 37°C 5 % CO2. Cells were pelleted for 5 min at 350 g. Trypsinized cells were then re-suspended in 7.5 mg/ml basement membrane matrix (Cultrex BME RGF type 2 (BME-2), Sigma Aldrich) supplemented with complete media and plated as 15 μl droplets in a 6-well plate. Once BME-2 polymerized, complete media was added and plates incubated at 37 °C 5 % CO2. Complete media: AdDMEM/F12 medium supplemented with HEPES (1×, Invitrogen), Glutamax (1×, Invitrogen), penicillin/streptomycin (1×, Invitrogen), B27 (1×, Invitrogen), Primocin (1 mg/ml, InvivoGen), N-acetyl-L-cysteine (1 mM, Sigma), recombinant Wnt3a protein (100 ng/mL, AMSBIO, AMS.rmW3aH-010), recombinant R-Spondin 1 protein (500 ng/m, AMSBIO, AMS.RS1-H4221), recombinant Noggin protein (0.1 μg/ml, Peprotech), epidermal growth factor (EGF, 50 ng/ml, Peprotech), fibroblast growth factor 10 (FGF10, 100 ng/ml, Peprotech), Nicotinamide (10 mM, Sigma), SB202190 (10 μM, Stem Cell Technologies), and A83-01 (0.5 μM, Tocris).

For organoids staining, culture medium was aspirated from the wells, washed with PBS and fixed in 2 % paraformaldehyde in PBS for 30 min. Samples were blocked for 1 h in blocking buffer (0.5 % bovine serum albumin, 0.25 % fish skin gelatin, 1 % Triton X-100 and 10% donkey serum) in PHEM buffer (60 mM PIPES, 25 mM HEPES, 10 mM EGTA, and 4 mM MgSO4·7H2O). Tissues were incubated with primary antibodies (**Supplementary table 1**) overnight at RT using blocking buffer, followed by 4 washes with 0.2 % Tween-20 in PHEM buffer of a minimum 15 min. When indicated EdU incorporation was detected with Click-iT chemistry kit according to the manufacturer’s instructions (Life technologies, 23227). Next, whole-mounts or inserts were incubated overnight with 1 μg/ml DAPI (Sigma Aldrich, D9542) and secondary antibodies (1:500) in blocking buffer. When indicated Alexa fluor 647-wheat germ agglutinin (WGA, Invitrogen W32466) was added 1:200 and Alexa fluor 647 anti-human/mouse CD49f (Biolegend, 313610) was added 1:250. Afterwards, samples were washed 4×15 min with 0.2 % Tween-20 in PHEM buffer and mounted using Vectashield mounting media (Vector Laboratories, H-1000). Imaging was performed using a SP8 Leica confocal microscope with a 40 x objective with 1x digital zoom, optimal pinhole and line average, bidirectional scan, speed 400-600 hz, resolution 1024×1024

### Human esophageal epithelioid generation

Esophageal tissue was obtained from deceased organ donors. Written informed consent was obtained in all cases under ethically approved protocols (Research Ethics Committee references 15/EE/0152 NRES Committee East of England— Cambridge South and 15/EE/0218 NRES Committee East of England—Cambridge East). A segment of mid-esophagus was excised within 60 minutes of circulatory arrest and preserved in PBS until processing. Esophageal epithelium was peeled from the underlying muscle using forceps and most of the submucosa layer scraped using a scalpel. Then, the sample was cut in pieces (explants) which were placed on membrane inserts and cultured as described above for mouse esophageal epithelioid cultures. Immunostaining was performed as described above.

### RNA sequencing

RNA was obtained from epithelioids 1 week after medium change to mFAD or from mouse esophageal epithelium peeled and incubated with Dispase I (Roche catalog no. 04942086001) diluted at 1 mg/ml in PBS for 15 min. RNA was extracted using RNeasy Micro Kit (QIAGEN, 74106), following the manufacturer’s recommendations. Briefly, cells were washed with cold Hank’s Balanced Salt Solution-HBSS (Gibco, 14175-053) and lysis buffer was added to the insert. The integrity of RNA was analyzed by Qubit RNA Assay Kit (Invitrogen, Q32852). RNA-seq, libraries were prepared in an automated fashion using an Agilent Bravo robot with a KAPA Standard mRNA-Seq Kit (KAPA BIOSYSTEMS). In house adaptors were ligated to 100-300 bp fragments of dsDNA. All samples were subjected to 10 PCR cycles using sanger_168 tag set of primers and paired-end sequencing was performed on Illumina HiSeq 2500 with 75 bp read length. Reads were mapped using STAR 2.5.3a, the alignment files were sorted and duplicate-marked using Biobambam2 2.0.54, and the read summarization performed by the htseq-count script from version 0.6.1p1 of the HTSeq framework ^60,61^. Raw counts were normalized using the median of ratios method^62^. Markers for basal cell compartment, cell cycle, cell differentiation were selected from published scRNAseq data^34^. Markers of the different esophageal differentiation steps (**Supplementary Table 1**) were selected from scRNAseq data^42^.

### EdU proliferation and lineage tracing

For *in vivo* proliferation analysis, 10 μg of EdU in PBS were administered by intraperitoneal injection 1 h before culling. For *in vitro* proliferation analysis, epithelioids were incubated with 10 µM EdU for 1 h. For EdU *in vitro* lineage tracing, cells were incubated with EdU for 1 h, washed and kept in mFAD for an additional 48h. Esophageal epithelium wholemounts and cultures were fixed and stained with EdU-Click-iT kit and immunofluorescence as previously explained. EdU-positive basal cells were quantified from a minimum of 10 z-stack images using Fiji ImageJ.

### Copy number analysis

DNA extraction was performed using the QIAMP DNA microkit (Qiagen, 56304) following the manufacturer’s instructions. DNA from the ears of the same mice was extracted with the same method and used as germline controls. Whole Genome Sequencing at low coverage was performed on either HiSeq 4000 machine (Illumina) to generate 150-bp paired-end reads. A modified version of QDNAseq (https://github.com/ccagc/QDNAseq/) was used to call changes in total copy number from the low coverage whole genome sequencing data^63^. QDNAseq was modified to include the correction of the coverage profile of the sample of interest by that of a matched control. Briefly, the procedure to call gains and losses is as follows: First sequencing reads were counted per 100 kb bins for both the sample of interest and the matched control. The bin-counts were then combined into coverage log ratio values to obtain what is commonly referred to as “logR”. The calculation of logr is implemented similarly to how the Battenberg copy number caller calculates these values^64^: first the bin-counts from the sample of interest were divided by the control bin-counts to obtain the coverage ratio; the coverage ratio was then divided by the mean coverage ratio and finally the log2 was taken to obtain logr. The standard QDNAseq pipeline is then continued with first a correction of the logr for GC content correlated wave artefacts, segmentation and finally calling of gains and losses.

A post-hoc filtering step was subsequently applied to obtain robust copy number calls. We noticed that several regions were commonly called as altered due to coverage local inconsistencies in the matched controls. To identify the effect of these inconsistencies we applied the copy number pipeline in a run where each control was matched against all controls from the whole genome sequencing data described in ^27^, expecting no alterations to be called. The run revealed common regions of false positive alterations on a number of chromosomes. Regions that were called in 3 or more different control-vs-control runs were subsequently masked from any analysis, i.e. copy number calls in these regions were not accepted.

Finally, after applying the masking we further filtered calls requiring a gain or loss to span at least 30Mb in size and that the alteration must constitute a gain or loss in at least 20 % of cells. To obtain an estimate of the percentage of sequenced cells that contained an alteration we applied the procedure that copy number caller ASCAT uses to find a tumor purity value^65^. ASCAT uses a grid search step where a range of purity and ploidy combinations are considered and ultimately a combination is picked by optimizing the amount of the genome that can be fit with an integer copy number value. We used this approach to estimate purity values only by fixing the ploidy at 2 and optimizing across a range of possible purity values.

The pipeline code and modified QDNAseq package are available at https://github.com/sdentro/qdnaseq_pipeline and https://github.com/sdentro/QDNAseq/tree/dev_respectively.

### Adenoviral infection

Cells from mice bearing inducible alleles were infected with Null adenovirus (Ad-CMV-Null, Vectorbiolabs, #1300) or *Cre*-expressing adenovirus (Ad-CMV-iCre, Vectorbiolabs, #1045), incubating them in 0.5 ml cFAD with 3.75×10^7 PFU/ml supplemented with 4 μg/ml Polybrene (Sigma Aldrich, H9268) on top of the insert and 2 ml cFAD on the bottom, at 37 °C, 5 % CO_2_ for 24 h. Next, cells were washed and fresh medium was added. Infection rates were > 90 %.

### SCA lineage tracing

Esophageal epithelioids from *Rosa26*^*confetti/confetti*^ animals were induced *in vitro* using adenovirus-Cre as specified above. A week after induction, medium was changed to mFAD and whole wells were imaged in an Incucyte system as specified before. Images obtained were analyzed using Fiji ImageJ.

Individual SCA that were recognizable at week 4 of imaging were followed until week 19 or until its small size makes it undistinguishable from the background. Areas were quantified at the specified time points. The average area from week 4 to 8 (A4-8), week 10 to 13(A10-13) and week 15 to 19 (A15-19) was used to classify the pattern of trajectory of each clone. SCA with A10-13=0 and A15-19=0 were classified as “Decay1”, SCA with A10-13>0 and A15-19=0 were classified as “Decay2”, other SCA with A4-8>A10-13>A15-19 were classified as “Decay3”. SCA with A4-8<A10-13>A15-19 were classified as “Biphasic”, SCA with A4-8<A10-13<A15-19 were classified as “Growing”, the rest of areas were classified as “Steady”. Wilcoxon matched-pairs signed rank test was used between the week 4, week 11 and week 19 were used to confirm the trajectories of each pattern.

### Quantitative SCA analysis and theoretical modelling

A least-squares minimization procedure was used to simultaneously fit the average area of SCA and the number of labelled SCA over time according to a single, equipotent progenitor model that describes proliferating cell behavior in esophageal epithelium *in vivo*^11,45^. The growth of the average SCA area was fitted to the theoretical linear expectation, i.e. a model of type *a*_0_ + *bt*, while the decline in the total fraction of labelled SCA was described by an hyperbolic function of type *N*_0_/(1 + *λ*′*t*), with constrained parameters (*λ*^*′*^ = *b*/*a*_0_, according to theory)^45^. Optimum parameter values 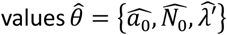 were obtained by averaging goodness-of-fit values, measured as the sum of the squared residuals (*y*_*i,obs*_ − *y*_*i,model*_) relative to the standard deviation of the observable (*σ*_*yi,obs*_), in both datasets. A zero-parameter fit followed for the total labelled area, modeled as constant *a*_0_*N*_0_ given by the product of the average SCA size and the total number of surviving SCA at different time points, which is consistent with homeostasis. The first two time points were ignored in the fits to avoid initial stabilization-related effects.

For simulations of clonal dynamics, a 2D lattice implementation of a stochastic Moran process was adopted, where (clonogenic) progenitor cells were set to compete neutrally in a 200×200 squared (*k*=8 neighbors, default) or hexagonal (*k*=6 neighbors) grid with periodic boundary conditions ^19,66^. A replacement rate *Λ* = 2*λ*′*φ* was selected to meet the inferred SP-model kinetic conditions, *φ* being a scaling factor (no. cells/grid unit) used for tractable clone simulations, a parameter that was later regressed out before readout. Simulation results are shown overlaid on experimental data, with shaded areas reflecting 95 % plausible intervals given by limited sample sizes equivalent to those in the experimental data (at least 200 permutation-built subsets).

The code developed for the quantitative clonal analysis has been made publicly available and can be found at https://github.com/gp10/ClonalDeriv3D.

### Single-colored area cutting and DNA sequencing

Single-colored areas were dissected using an LMD7 microscope (Leica Microsystems) and collected in separate tubes. Samples were digested and DNA extracted using the QIAMP DNA microkit (Qiagen, 56304) following the manufacturer’s instructions. DNA from the ears of the same mice was extracted with the same method and used as germline controls.

DNA sequencing was performed using a custom bait capture of 192 frequently mutated genes in cancer as in^19^, briefly samples were multiplexed and then sequenced using an Illumina HiSeq 2500 and paired-end 75-base pair (bp) reads. Alignment was performed using BWA-MEM (v.0.7.17, https://github.com/lh3/bwa)52 with optical and PCR duplicates marked using Biobambam2 (v.2.0.86, https://gitlab.com/german.tischler/biobambam2, https://www.sanger.ac.uk/science/tools/biobambam). The mean coverage was 106.2 x, ranging from 69.57-157.5 times between SCA.

### Generation of p53* mutant transformed cells (p53*-TC)

Ahcre^ERT^-*Trp53*^*flR245W-GFP/wt*^ mice were induced to express the mutant *p53*^*R245W*^ allele and GFP reporter protein, by intraperitoneal injection of 80 mg/kg β-naphthoflavone (MP Biomedicals, 156738) and 0.25 mg Tamoxifen (Sigma Aldrich N3633) as previously described^28,30^. Once month later, mice were orally treated with the carcinogen diethylnitrosamine (DEN) (Sigma, catalog no. N0756) in sweetened drinking water (40 mg per 1,000 ml) for 24 h on 3 days a week (Monday, Wednesday and Friday), for two weeks to induce the formation of early lesions in the esophageal epithelium, followed by exposure to 10 doses of 2Gy of ionizing radiation using a whole-body caesium source irradiator. Mice were sacrificed and tumors were collected and cultured. After several rounds of expansion, cells were assessed for GFP expression (100% of the culture).

p53*-TC were subjected to M-FISH using mouse painting probe by the Molecular Cytogenetics Core Facility at Wellcome Sanger Institute. Briefly, 20 randomly selected metaphases were karyotyped based on M-FISH and DAPI-banding patterns. The results were analyzed focusing on karyotype instability and heterogeneity in terms of structural and numerical aberrations.

### Flow cytometry

Keratinocyte cultures were detached by incubation with 0.05 % Trypsin-EDTA for 20 min at 37°C 5 % CO_2_. Cells were pelleted for 5 min at 650 g and resuspended in PBS to be immediately analyzed using a Becton Dickinson (BD) LSRFortessa. Where suprabasal and basal cells needed to be quantified, cells were fixed using 2 % PFA and incubated in blocking solution (0.1 % BSA 0.5 mM EDTA in PBS) for 15 min, followed by an incubation with anti-ITGA6-647 antibody (1:125, Biolegend, UK 313610) in blocking solution for 45 min at RT. YFP fluorescence was collected using the 488 nm laser and the 530/30 bandpass filter and ITGA6-647 fluorescence was collected using the 640 nm laser and the 670/14 bandpass filter. Data was analyzed using FlowJo software (version 10.5.3). Basal cells were defined as ITGA6 positive cells and suprabasal cells as ITGA6 negative cells.

### Cell competition assays

The indicated cell populations were trypsinized and mixed, 1:1. After one week in cFAD, when the cultures are fully confluent, medium is changed to mFAD and starting time point was collected. At the time points specified, cells were collected and analyzed by flow cytometry or fixed for microscopy as stated before. Where indicated cells were treated with 2 hour pulses of epirubicin 1µM, 5-fluorouracyl 5µM twice a week or irradiated with 2Gy (once a week using an Xstrahl CIX2 or RPS CellRad RSM-009 irradiators). Cell fitness over YFP^+^ or nTnG cells was measured by quantifying fold increase of YFP-or p53*-TC cells respectively at the specified time point versus its proportion at the initial time point.

## DATA AVAILABILITY

The sequencing data sets in this study are publicly available at the European Nucleotide archive (ENA) Accession numbers for RNAseq data on https://www.ebi.ac.uk/ena are as follows:

*In vivo* samples: ERS14340821, ERS14340822, ERS14340823, ERS14340824. *In vitro* samples: ERS2515249, ERS2515250, ERS2515251, ERS2515252.

Accession numbers for targeted DNA sequencing of SCA is ERP107379.

Data used to generate each figure is available in **Supplementary table 1**.

---

**Extended Data Fig. 1:**
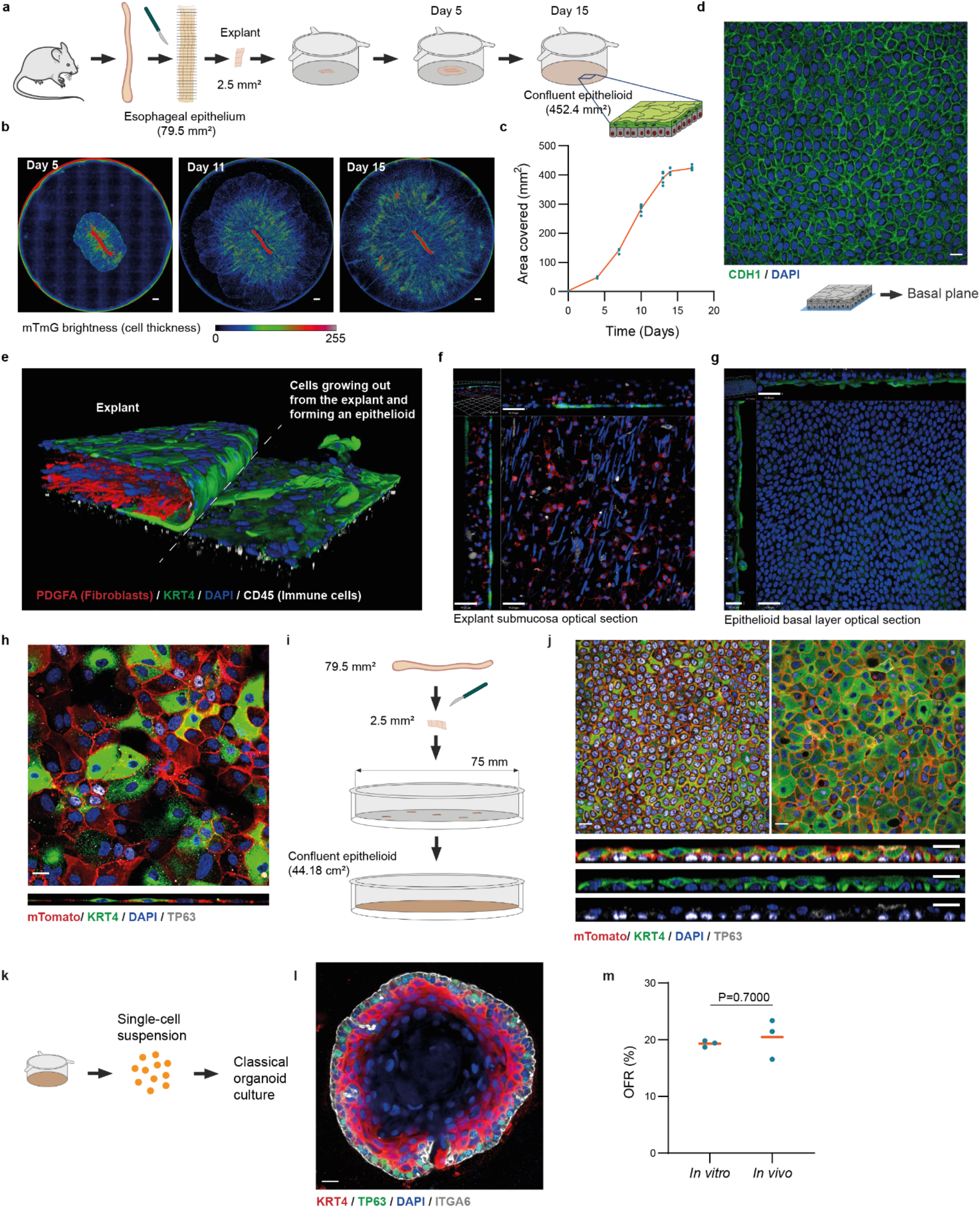
Generation and amplification of mouse esophageal epithelioids. **a**, Schematic illustration of epithelioid generation from a single esophageal explant. **b**, Live microscopy ‘Incucyte’ images 5, 11 and 15 days after plating *Rosa26*^*mTmG*^ esophageal explants. mTomato fluorescence is shown using the Rainbow LUT of Fiji ImageJ. Scale bar, 1000 μm. **c**, Area of culture insert covered by cells over time from a single esophageal explant. Each dot represents an epithelioid. *n=* 6 epithelioids. **d**, Optical section of basal layer of epithelioid stained for CDH1 (green) and DAPI (blue). Scale bar, 20 μm. **e-g**, Immunostaining against PDGFA (red, fibroblasts), KRT4 (green), CD45 (grey, immune cells) and DAPI (blue), of an explant and the cells exiting from it to colonize the insert and form an epithelioid. (e) 3D image (e) and optical section with orthogonal views of the submucosa layer of the explant (f, scale bars=44 μm) and its newly formed epithelioid basal layer membrane (g, scale bars=41 μm). **h**, Primary *Rosa26*^*mTmG*^ mouse esophageal keratinocytes cultured in a plastic surface stained for TP63 (grey), KRT4 (green) and DAPI (blue). Basal layer section (top panel) and lateral view (bottom panel). Scale bar, 20 μm. **i-j**, 5 explants from *Rosa26*^*mTmG*^ mouse esophagus plated in a 75mm diameter insert and grown for 20 days. (i) Plating scheme (j) and optical sections of basal layer (top-left), suprabasal layer (top-right) and lateral views (bottom) of a representative area of the culture stained for TP63 (grey), KRT4 (green) and DAPI (blue). Scale bars, 20 μm. **k-m**, Organoid generation from epithelioids. Protocol (k), representative image of esophageal organoid (l) stained for ITGA6 (grey), KRT4 (red), Ki67 (green) and DAPI (blue), scale bar=14 μm, and organoid formation rate (OFR, %) (m) from esophageal epithelioids and mouse esophagus. *n=*3 replicates from different original epithelioid cultures or mice. Orange bars indicate mean values. Two-tailed Mann-Whitney test. Source data in Supplementary Table 1.

**Extended Data Fig. 2:**
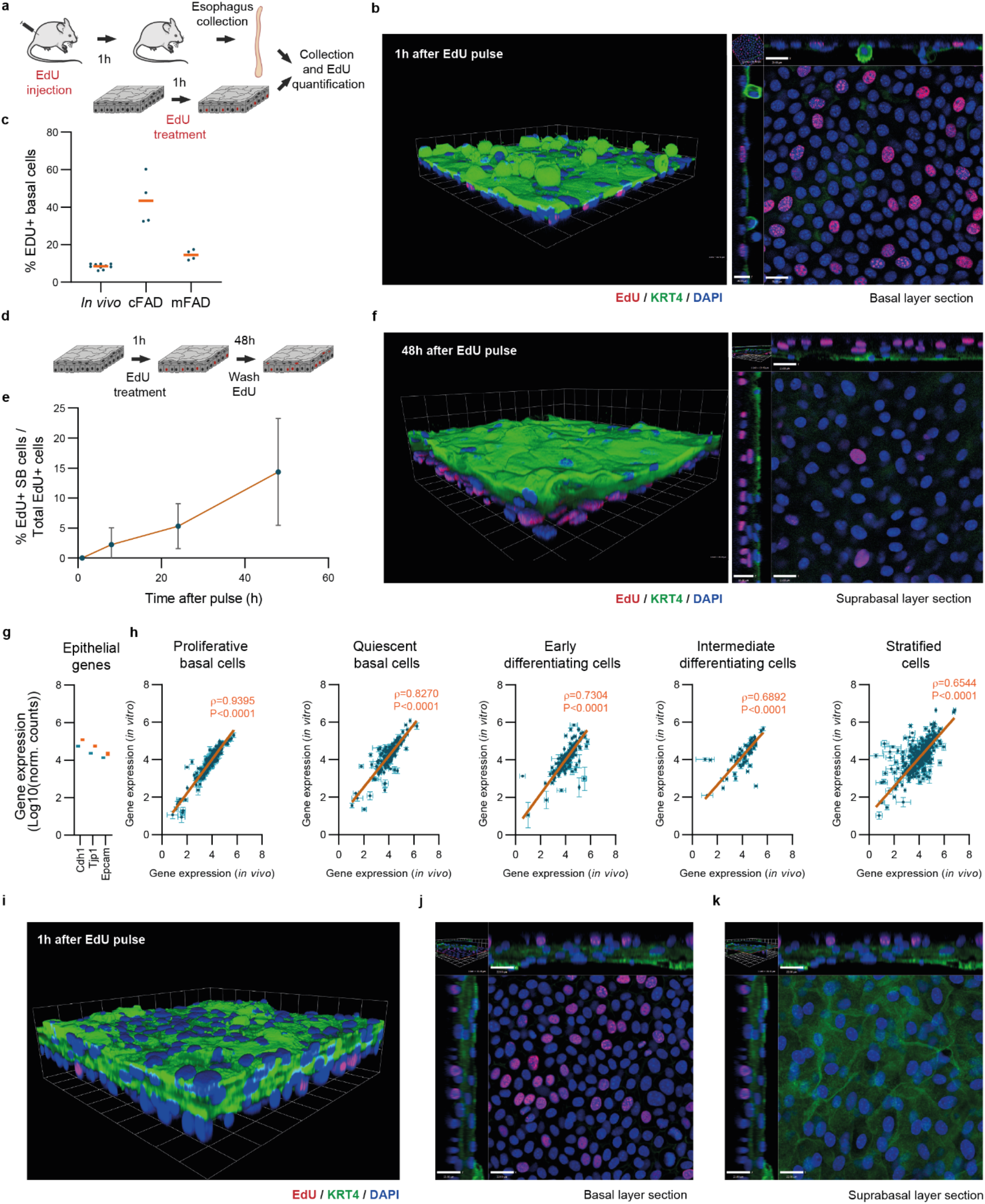
Characterization of mouse esophageal epithelioids. **a-c**, Proliferating cells that incorporated EdU after a 1h pulse in mouse esophageal epithelium and mouse esophageal epithelioids treated in minimal medium (mFAD) or complete medium (cFAD). **a**, protocol. **b** rendered confocal z stack (left) and basal plane section with orthogonal views (right) of a typical mouse esophageal epithelioid in mFAD stained for KRT4 (green), EdU(red) and DAPI (blue). Scale bars=20 μm for x-y and 14 μm for z plane. **c**, Percentage of EdU^+^ basal cells in the three conditions, each dot corresponds to an animal or an epithelioid, orange bars represent the mean values. **d-f**, *In vitro* proliferative cell tracing 48h after an EdU pulse. **d**, Protocol, EdU labels S phase cells, after 48 hours EdU^+^ cells that have differentiated and left the basal layer are counted, revealing the rate of differentiation and stratification **e**, percentage of EdU^+^ suprabasal cells versus total EdU^+^ cells (mean±SD). **f**, rendered confocal z stack (left) and basal plane section with orthogonal views (right) of typical mouse esophageal epithelioid stained for KRT4 (green), EdU (red) and DAPI (blue). Scale bars=21 μm for x-y and 18 μm for z plane. **g-h** RNA sequencing comparing gene expression from mouse esophageal epithelium (*in vivo*) and esophageal epithelioids (*in vitro*). *n=*4 animals and 4 epithelioids from 4 different animals. **g** Epithelial genes and correlation between *in vivo* and *in vitro* samples in transcripts characterizing the different stages of differentiation^42^. The orange line shows the linear regression between samples with the Pearson’s coefficient and p-value of the correlation. **i-k**, Rendered confocal z stack (**i**), basal plane section with orthogonal views (**j**), suprabasal plane section with orthogonal views (**k**) of a typical mouse esophagus epithelioid exposed to air-liquid interface for 15 days, incubated for 1h with EdU and stained for KRT4 (green), EdU (red) and DAPI (blue). Scale bars=22 μm for x-y and 22 μm for z plane. Source data in Supplementary Table 1.

**Extended Data Fig. 3:**
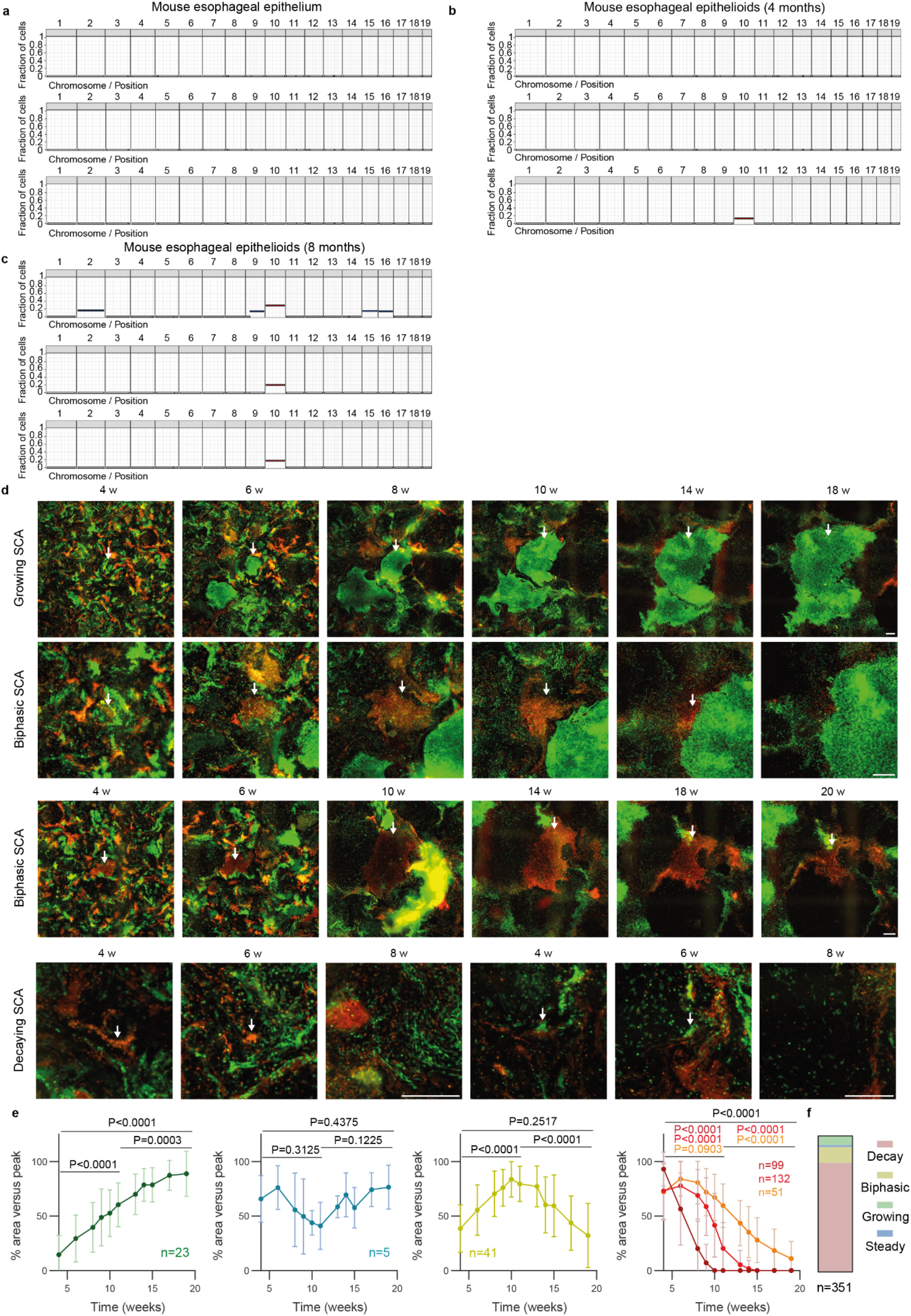
Long-term maintenance and tissue dynamics of esophageal epithelioids. **a-d**, Whole genome sequencing of mouse esophageal epithelium and epithelioids after 4 and 8 months in culture. *n=*3 animals and 3 esophageal epithelioids from different animals per time point. Graphs showing the fraction of cells bearing a DNA gain or loss in each chromosome in normal esophageal epithelium of three mice (events present in more than 10 % of cells) (**a**), epithelioids from three different mice after 4 (**b**) and 8 (**c**) months in culture. **d-e**, Epithelioids from *Rosa26*^*confetti/confetti*^ mice after *in vitro* labelling (see methods), cultured for 24 weeks. Representative examples (**d**) and area quantification (**e**) of 351 SCA showing growing, biphasic, constant and decreasing changes in area. Arrows indicate selected SCA. Wilcoxon matched-pairs signed rank test. Mean±SD per time point and pattern are indicated. Areas of SCA, grouped by change in area (see methods), as indicated in each graph. **f**, Proportion of SCA showing each group. Source data in Supplementary Table 1.

**Extended Data Fig. 4:**
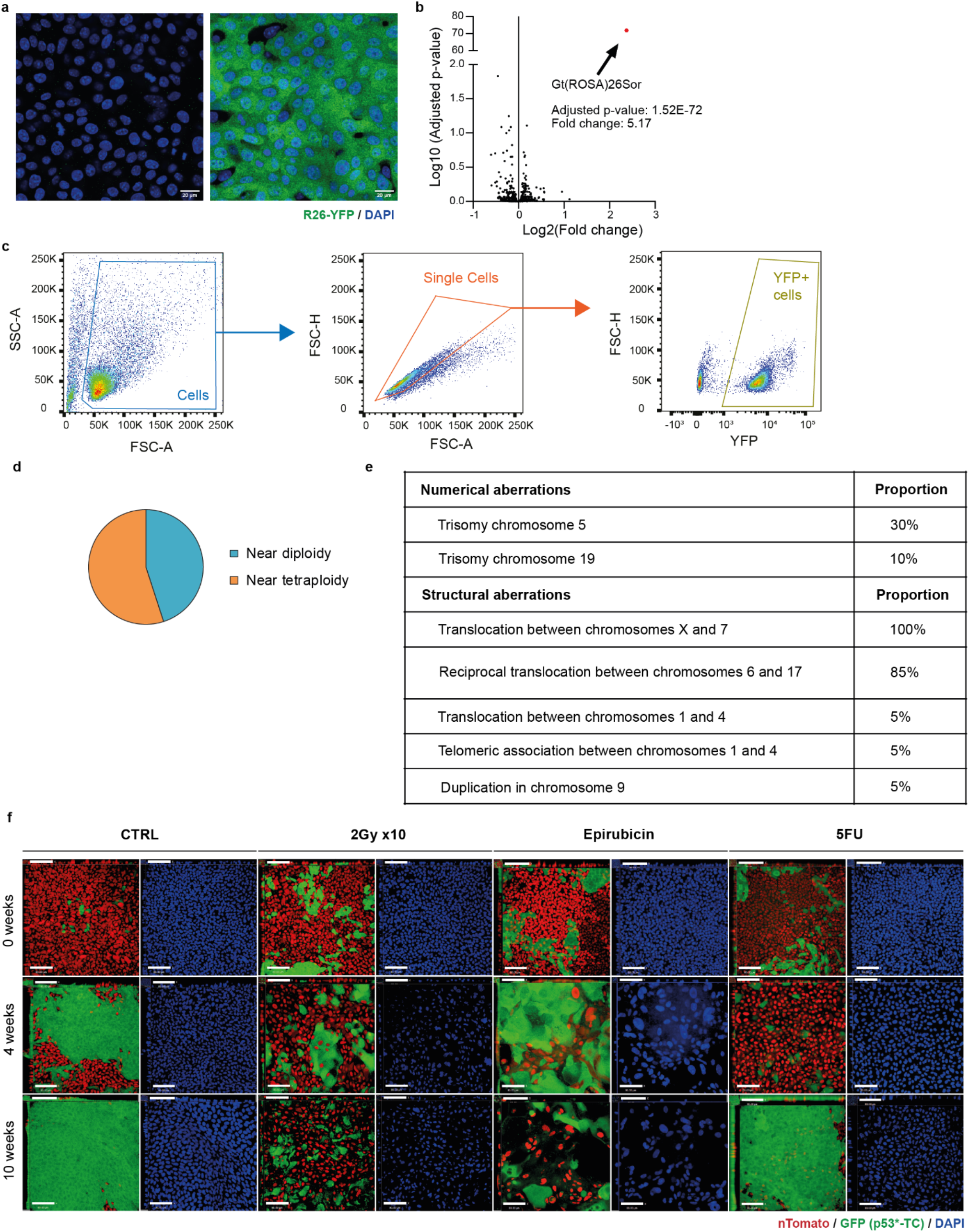
Epithelioids as a tool to study cell competition. **a**, Uninduced (left, YFP-) or induced (right, YFP^+^) cells from mouse esophageal epithelioids from *Rosa26*^*YFP/YFP*^ mice. Scale bar, 20 μm. **b**, Volcano plot comparing RNA expression (Log2 of the fold change) of induced and uninduced cells from **a** and log10 of adjusted p-value. Red dot shows the only significant gene. Wald test corrected for multiple testing using the Benjamini and Hochberg method. **c**, Gating strategy for quantifying the YFP^+^ cell population in cell competition experiments with YFP^+^ and non-fluorescent subpopulations. **d-e**, M-FISH karyotyping analysis of *p53*^*R245W*^ mutant cancer cells (p53*-TC) used in Fig.5. **e**, Images shown in **Fig.5 i-l** with their corresponding panels of DAPI channel in blue. Scale bars, 80 μm. Source data in **Supplementary Table 1**.

